# An integrated identifiability and metabolic flux analysis workflow for reducing and repairing dynamic CHO metabolic models

**DOI:** 10.64898/2026.10.26.761781

**Authors:** Shivam Barodiya, Nikola Malinov, Srikanth Rapala, Sarah Harcum, Marianthi Ierapetritou

**Affiliations:** University of Delaware; Clemson University

**Keywords:** Metabolic modeling, Perfusion, Identifiability analysis, Metabolic flux analysis, Amino acid supplementation

## Abstract

Over-parametrized kinetic-metabolic models can fit available data accurately yet still yield unreliable parameter estimates and output uncertainties. We developed an integrated workflow that combines Fisher information matrix-based identifiability analysis diagnostics, uncertainty propagation, and metabolic flux analysis (MFA) to reduce model size and assess the transferability of a kinetically constrained dynamic flux balance analysis (dFBA) model of CHO perfusion culture. Using Control medium data, the dFBA model size was reduced from 48 to 43 kinetic parameters while maintaining performance across 23 measured extracellular species. The reduced model’s transferability was assessed by predicting cell behavior in an amino acid-supplemented medium (Medium C), which lowered the specific growth rate while increasing specific productivity. The reduced model successfully predicted viable cell density, antibody titer, and supplemented amino acids for Medium C but failed to capture the reversal of glutamate metabolism and the increase in lactate production. Output uncertainty quantification identified structural gaps rather than parameter uncertainty as the underlying cause of prediction failures, with 79% of Medium C experimental points lying outside the 95% parameter-propagated interval. Stoichiometric MFA linked the Medium C phenotype to increased amino acid catabolism, identified the intracellular sources for glutamate and lactate production, and enabled estimation of two new rate-constants for their production directly from stoichiometric flux, rather than fitted to data. MFA-guided repair reduced glutamate and lactate NRMSE in Medium C by 64% and 36%, respectively. The workflow provides a systematic strategy for improving model identifiability and using MFA to diagnose and repair structural discrepancies.

**Author summary:** Computer models of cell biology are often complex, with more parameters than the data can support. Such models can explain the data well while still containing unreliable parameters and may fail on unseen data. We developed a workflow that identifies which model parameters are supported by the data and simplifies the model accordingly while maintaining accuracy. We applied this to a computational model of CHO cells, the cell type widely used to manufacture biopharmaceutical antibody drugs. After simplifying the model, we tested whether it still worked when the cells were grown in a different nutrient medium, and it failed to predict two key metabolites, glutamate and lactate. Rather than fixing the issue by adding more parameters, a common but not the best method, we used a complementary technique to pinpoint exactly which biological process was missing from the model. We calculated the missing values directly from measured data rather than fitting them to the data we were trying to predict. This repaired the model structure. Our workflow is a general strategy for building cell biology models that are reliable when applied beyond the exact conditions under which it was built.

## 1. Introduction

Mathematical models of cellular metabolism provide insights into its mechanisms (Kyriakopoulos, Ang et al. 2018, Tang, Xu et al. 2020). However, models can become very large depending on the extent of the cell metabolism captured. The model size is determined by the number of species and the number of model parameters. Kinetic models can range up to tens of species and parameters (Nolan and Lee 2011, Ben Yahia, Malphettes et al. 2015). Given the time- and resource-intensive nature of data collection, it is essential to use experimental data efficiently (Franceschini and Macchietto 2008, Waldron, Pankajakshan et al. 2019). Small-sized models require less data, making them easier to adapt and use for process optimization. With many kinetic parameters and limited datasets, it becomes difficult to confidently estimate all of the parameters (Fekih-Salem, Dewasme et al. 2019).

Identifiability analysis can expose redundant parametrization and quantify confidence in parameter estimates, helping to reduce model size. MFA can reveal metabolic rearrangement and intracellular origins of metabolic activities. However, these analyses are generally used separately rather than as part of a unified workflow to construct identifiable kinetic metabolic models that are transferable across conditions. Linear algebra-based mathematical tools have been developed and used to assess confidence in parameter estimates, and to quantify uncertainty in model outputs for identifiability analysis (Franceschini and Macchietto 2008, Wang and Dowling 2022). A kinetic model reduced and validated on one condition is often subsequently applied to a new condition without first verifying whether its structure remains valid. When structural discrepancies arise, traditional approaches often address these gaps by introducing additional fitted empirical parameters without the guidance of MFA. This practice can mask the underlying mechanistic gaps and inflate model size. Combining identifiability analysis with MFA prevents this issue; MFA can localize the gap in the model’s structure and supply the corresponding rate terms directly from measured flux data, circumventing the need to introduce additional fitted parameters.

This work integrates identifiability and MFA to reduce and repair the structure of a recently developed kinetically constrained dynamic flux balance analysis (dFBA) model (Reddy, Malinov et al. 2026). Fisher information matrix (FIM) based identifiability analysis was used to reduce model size while preserving model performance (Wang and Dowling 2022). Next, we assessed the model’s transferability using an amino acid-supplemented formulation that lowered the specific growth rate while increasing specific productivity, producing a distinct phenotype. Finally, uncertainty analysis and MFA were used to distinguish parameter uncertainty from structural failure and to repair the model without introducing additional fitted parameters. By localizing the metabolic origin of structural failure and deriving repair terms directly from flux data, this approach avoided incorporating each new phenotype as an additional parameter. The kinetically repaired reduced model structure was validated by testing the identifiability of model parameters on the combined dataset.

## 2. Materials and Methods

This section describes the dFBA model, details the experimental data, and outlines the mathematical tools used for model reduction, repair, and performance assessment.

### 2.1 Kinetically constrained dynamic flux balance analysis model

To study the metabolism of CHO-K1 VRC01 cells cultured in perfusion mode in the ambr250 system, the published dFBA model was applied (Fig 1) (Reddy, Malinov et al. 2026). The model integrates a kinetic component with a stoichiometric network and mass balances to calculate the concentrations of extracellular measured species for all experimental time points. An extensive dataset was collected to train and assess the dFBA model, including the measurements of glucose, lactate, ammonia, 18 canonical amino acids, cell density, and antibody titer, a total of 23 measurements for all experiments.

**Fig 1.**
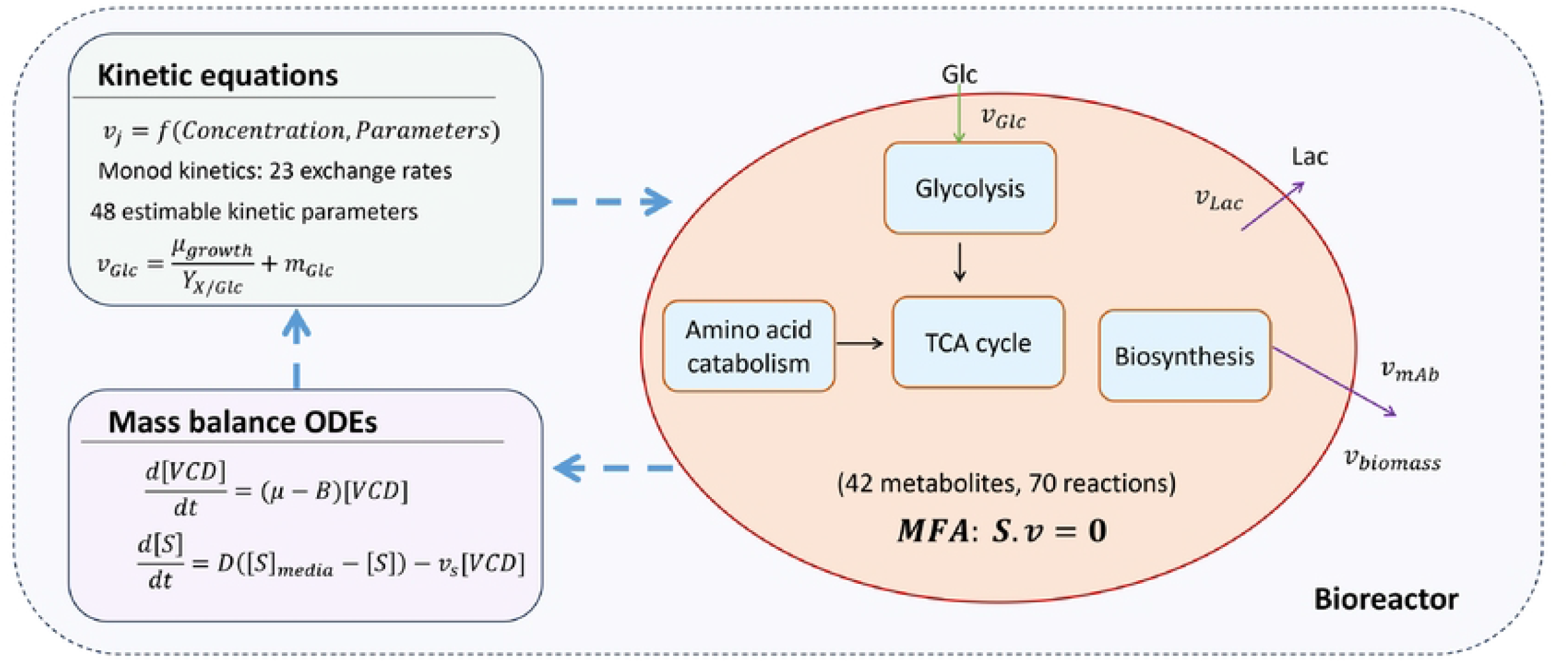
Schematic of the dFBA model, integrating kinetic framework and stoichiometric network to simulate intracellular metabolism and perfusion bioreactor dynamics over time.

The initial model contained 48 kinetic parameters across all rate equations (Table A1). The stoichiometric component had 42 metabolites and 70 reactions to explain cell metabolism. As compared to the published model, 2 reactions were added to the network, arginine to glutamate and glycine to ammonia, and the biomass reaction was modified to capture the cell metabolism (Table B1) (Reddy, Malinov et al. 2026). To obtain the intracellular species fluxes and constrain the calculated extracellular fluxes, MFA was applied at each experimental time point. Based on the reaction network, the stoichiometric component provided biologically relevant bounds on the extracellular species fluxes while solving for intracellular species fluxes.

The MFA formulation used here solves a weighted least-squares loss between calculated and kinetically predicted exchange fluxes, subject to the pseudo-steady-state mass balance (Sv = 0, where S is the stoichiometric flux and v is the flux vector), with no assumed cellular objective. This differs from the kinetically constrained dFBA formulation of the base model (Reddy, Malinov et al. 2026), which combined the measurement-fit terms with an explicit flux-parsimony objective.

The kinetic component contained Monod equations to calculate the exchange flux for extracellular species, with extracellular species concentrations and initial kinetic parameter estimates as inputs. The extracellular species flux output of the kinetic model served as an input to solve the stoichiometric network.

After obtaining the flux values at a single time point using the stoichiometric network, discretized mass balances were applied to the bioreactor and viable cells to update the extracellular species concentrations for the next time point. Updated concentrations served as input to the kinetic component, allowing the model to be solved for all time points in the experiment. Kinetic model parameters were estimated using the dual annealing method for global parameter search, and the dual active set quadratic programming (DAQP) solver was deployed to obtain intracellular fluxes from MFA.

### 2.2 Dataset for identifiability analysis, uncertainty quantification and structural repair

CHO-K1 cell line expressing an anti-HIV monoclonal antibody (VRC01) was cultured in perfusion mode in ambr250 bioreactors. Dynamic perfusion and steady-state operations were performed using two different perfusion media, called Control and Medium C (Table 1). In the first experiment, the ambr250 system was operated in a dynamic perfusion mode (without cell bleed) to achieve the maximum cell density for the Control medium formulation (Table 2). In the next experiment, the bioreactor was operated at two consecutive steady states of 40 × 10^6^ and 25 × 10^6^ cells/mL, with the same Control medium formulation as in the dynamic perfusion condition. The in-house perfusion medium, composed of HyClone Actipro basal medium and Cell Boost, constituted the Control medium (Malinov, Barodiya et al. 2026). Collectively, the dynamic perfusion condition and the two sequential steady states (Control SS40 and Control SS20) using the same Control medium formulation are referred to as the Control condition. All three Control condition datasets (dynamic perfusion, Control SS20, and Control SS40) were used collectively for mode reduction by combining and fixing kinetic parameters while retaining model performance. Subsequent comparisons to Medium C use only matched steady-state pairs – Control SS20 with Medium C SS20, and Control SS40 with Medium C SS40.

**Table 1.** Perfusion medium composition for Control and Medium C.

| Medium | Glucose | HyClone ActiPro basal medium | HyClone Cell Boost 7a | HyClone Cell Boost 7b | Arginine and leucine supplementation |
| --- | --- | --- | --- | --- | --- |
| Control | 50 mM | Yes | 3% (v/v) | 0.3% (v/v) | No supplementation |
| Medium C | 50 mM | Same as Control | Not added | 1.5% (v/v) | Arginine medium concentration: 5mM<br>Leucine medium concentration: 12 mM |

**Table 2.** Experimental data description with two perfusion media, Control and Medium C. Each condition had two biological replicates and a perfusion rate of 1 vessel volume per day (*vvd*^−1^)

| Condition | Operating mode | Cell density set-point | Medium | Used for |
| --- | --- | --- | --- | --- |
| Control dynamic perfusion (no cell bleed) | No bleed | Maximum achievable | Control | Model size reduction |
| Control SS20 | Steady state | $25 \times 10^6$ cells/mL | Control | Model size reduction |
| Control SS40 | Steady state | $40 \times 10^6$ cells/mL | Control | Model size reduction |
| Medium C SS20 | Steady state | $20 \times 10^6$ cells/mL | Medium C | Transferability test |
| Medium C SS40 | Steady state | $30 \times 10^6$ cells/mL | Medium C | Transferability test |

To assess the transferability of the reduced identifiable model obtained using the Control condition, data from Medium C were used. In the Control dynamic perfusion condition, a subset of amino acids was depleted at high cell densities, and these were supplemented in Medium C to increase cell-specific productivity while reducing the specific growth rate. Medium C was a metabolic perturbation designed to produce a phenotype distinct from that of the Control medium.

In Medium C, the total amino acid pool was reduced by omitting HyClone Cell Boost 7a, which provided additional amino acids to enhance cellular performance. Four essential amino acids, tryptophan, tyrosine, arginine, and leucine, were supplemented to increase the ratio of essential to non-essential amino acids in perfusion Medium C (Xing, Kenty et al. 2011). Tyrosine and tryptophan were supplied by increasing the volume fraction of HyClone Cell Boost 7b to 1.5% (v/v) relative to the Control medium (0.3% v/v) (Table 1). Due to a lower amino acid pool, the Medium C ambr250 system was first operated at a cell density set point of 20 × 10^6^ cells/mL and then at 30 × 10^6^ cells/mL. To compare the cell metabolism of Medium C with Control medium, cell density states (25 × 10^6^ and 40 × 10^6^ cells/mL) for the Control medium are compared with (20 × 10^6^ and 30 × 10^6^ cells/mL), respectively, for Medium C. Medium C provided a drastically different cell metabolism than Control medium when operated at similar cell density steady states.

### 2.3 Identifiability-guided model reduction and structural repair framework

Starting with the initial model, it was evaluated for over-parametrization and non-identifiable parameters using FIM-based identifiability analysis. Eigenvalues and eigenvectors were used to determine non-identifiable parameters using the Control condition. Identifiability analysis investigated mathematical structure of rate equations and confidence in parameter estimates. If certain parameters were non-estimable, they were either combined or replaced to improve model identifiability, thereby reducing model size (Franceschini and Macchietto 2008).

To assess the transferability of the reduced identifiable model, it was challenged to predict the cell metabolism under Medium C, where it failed to predict some species. The uncertainty analysis on the reduced model helped distinguish model parameter identifiability from structural discrepancy and identified structural gaps as the source for prediction failure. MFA-guided diagnosis was used to investigate rate equations and to repair them based on cell metabolism (Antoniewicz 2015). The repair prevented the addition of data-driven estimates of new kinetic parameters and any further model-size expansion, resulting in a reliable, transferable model with identifiable parameters and predictive capabilities. A general overview of the workflow is presented in Fig 2.

**Fig 2.**
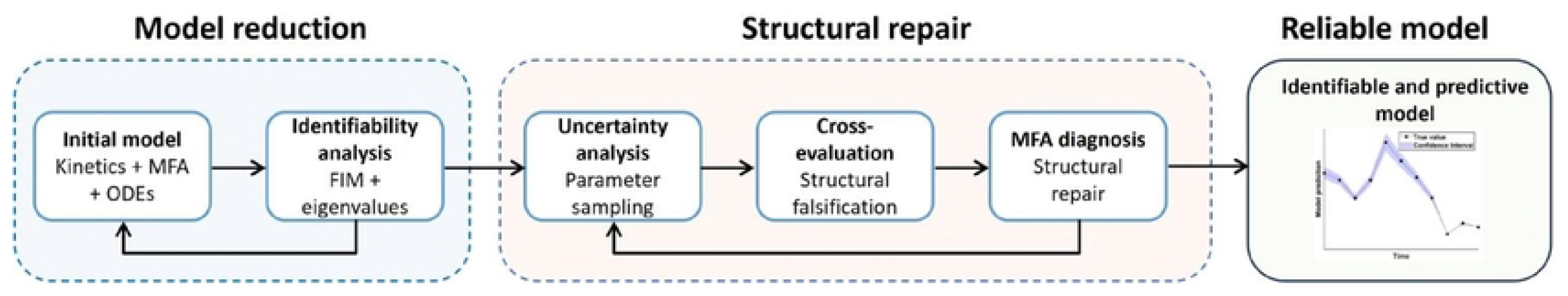
Identifiability-guided model reduction and MFA-guided structural repair workflow.

For the identifiability analysis, FIM-based eigenvalues and eigenvectors were used, which required computing sensitivities and FIM. The dynamic sensitivity of model outputs to model parameters helped determine their dependence on parameters. It was defined as derivatives of model outputs with respect to parameters, evaluated at every experimental time point for all the outputs and parameters (Equation 1) (Brun, Reichert et al. 2001, Franceschini and Macchietto 2008).

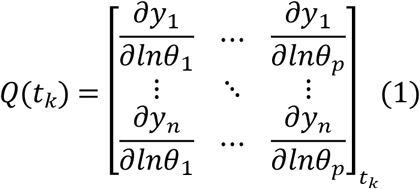

where y is the model output, *θ* denotes the model parameters, p is the number of parameters, and n is the number of measured outputs evaluated at each experimental time point *t*_*k*_. In this work, a central-difference method with 10% perturbation of each parameter was used to calculate the partial derivatives for log-space sensitivities (Equation 2) (Wang and Dowling 2022). Sensitivities were computed in log-parameter space because parameters spanned several orders of magnitude (e.g., 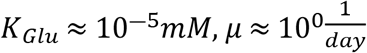). Working in log-space ensured that each parameter contributed comparably to the optimization landscape and the resulting FIM.

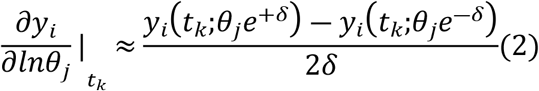

Where *i* ∈ {1,*n*}, *j* ∈ {1,*p*} and *δ* = ln (1.1) (10% multiplicative perturbation for each parameter).

FIM quantified the uncertainty in model parameters using dynamic local sensitivity and experimental noise, calculated as the time-summed, noise-weighted sensitivity product (Equation 3) (Franceschini and Macchietto 2008, Wang and Dowling 2022).

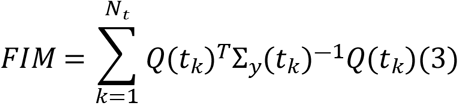

where Σ_*y*_(*t*_*k*_) is the noise covariance at time *t*_*k*_, *Q*(*t*_*k*_) is dynamic sensitivity at time *t*_*k*_, and the FIM summation runs over all *N*_*t*_ experimental time points.

All the identifiability computations were performed in log-parameter space to enforce parameter positivity. Log-parameter space spans several orders of magnitude in kinetic parameters and yields multiplicative confidence intervals (Raue, Schilling et al. 2013). Relative magnitudes of entries in *FIM*^−1^ were used to test confidence in model parameter estimates. The diagonal entries of *FIM*^−1^ were used to compute variance in model parameters, while off-diagonal entries were used to obtain pairwise covariance in model parameters. The FIM, being based on local sensitivity analysis, holds asymptotically true for non-linear models and is largely influenced by the initial parameter estimates (Raue, Kreutz et al. 2009).

Experimental noise captures the uncertainty in data arising from measurements. The FIM weighted each measurement by its inverse variance, requiring an estimate of the measurement noise covariance Σ_*y*_. With only two biological replicates per condition, a direct per time-point variance estimate could vanish when replicates coincided, making Σ_*y*_ singular with artificially low experimental noise. To obtain a smooth, strictly positive variance estimate, a heteroscedastic power-law variance model (Equation 4) was fitted to the replicate data for each measured species independently (Davidian and Carroll 1987, Franceschini and Macchietto 2008).

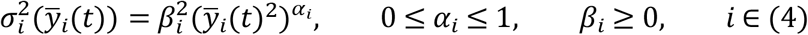

where 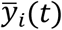 is the replicate mean of species i at sampling time t, and 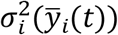 is the corresponding pooled sample variance across the two biological replicates. At each sampling time, 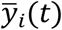, and 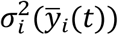 were computed from the two replicates data, and the parameters (*β*_*i*_, *α*_*i*_) were estimated for each species by least-squares fitting of Equation 4 to full set of (mean, variance) pairs pooled across all sampling times. Measurements were assumed uncorrelated across species and independent in time, a widely used assumption, giving a diagonal, time-dependent covariance matrix in Equation 5, which was utilized in Equation 3 for FIM calculations (Franceschini and Macchietto 2008).

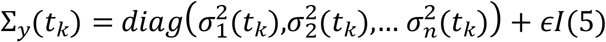

where a small regularization *ϵ* = 0.1 was used to ensure invertibility of the matrix.

Identifiability analysis represents confidence in estimating model parameters based on the model structure and available data (Raue, Kreutz et al. 2009). Parameters with an eigenvector magnitude ≥ 0.3 in each eigenvalue direction were considered dominant in that direction. This eigenvector threshold was a practical criterion following the common practice of using a value well above the uniform-contribution baseline (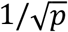 for p parameters). Parameters lying in eigenvalue directions with an eigenvalue magnitude close to zero were considered non-identifiable. Using the spectrum of eigenvalues for all parameter directions, the identifiability of parameters was determined.

The FIM rank (maximum number of identifiable parameters) and condition number (ratio of largest to smallest eigenvalue) were used to assess overall identifiability; their interpretation is provided in Supplementary file 1. The coefficient of variation for each parameter (*CV*_*j*_) was used to quantify the uncertainty in model parameters relative to their estimated value, as given by Equation 6. Standard deviation for each model parameter 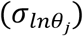 was obtained using diagonal entries in *FIM*^−1^ from Equation 7 (Fekih-Salem, Dewasme et al. 2019).

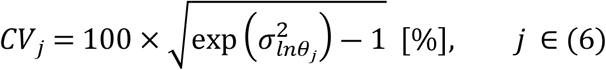

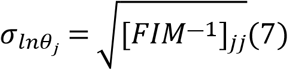

A pairwise correlation coefficient (*ρ*_*ij*_) (Equation 8) was calculated for the kinetic parameters to test their identifiability and mutual dependency (Fekih-Salem, Dewasme et al. 2019).

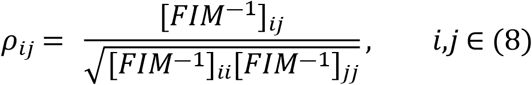

The asymmetric confidence interval for the model parameters was calculated using Equation 9 as reported in Supplementary file 2.

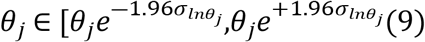

With the *FIM*^−1^ providing model parameter uncertainties, the uncertainties in model outputs were quantified by Monte Carlo parameter sampling. Parameter vectors were drawn from a multivariate normal distribution in log-parameter space, using the estimated parameter values as the mean and the full log-space covariance matrix *FIM*^−1^ as the covariance (Anane, López C et al. 2019). Sampling from full covariance preserved the inter-parameter correlation structure captured by FIM. Each of the 300 sampled parameter sets was propagated through the model to generate parameter-propagated uncertainties in model outputs. Based on the Cramer-Rao bound, *FIM*^−1^ represents the smallest possible parameter uncertainties, so the resulting parameter-propagated uncertainties were a lower bound, and the true uncertainty might be larger (Walter and Pronzato 1997).

The identifiability analysis highlighted structural inadequacies in the kinetic model for Medium C, which were biologically repaired using the MFA. MFA was applied to compare metabolic rewiring between Control and Medium C. For the flux-reversal species between Control and Medium C, MFA-guided selection of intracellular production sources, which was based on reactions showing more than 100% change in flux magnitude between the two media compositions.

To quantify the model performance improvement, the normalized root mean square error (NRMSE) and mean absolute error (MAE) were calculated, with the mean of the experimental data across all time points used as the normalization factor for RMSE (Bayer, Duerkop et al. 2023). NRMSE and MAE values highlighted improvements in the model following MFA-based structural repair.

## 3. Results

For the dFBA model, the kinetic component structure was evaluated using identifiability analysis. Eigenvalues and eigenvectors highlighted poorly identifiable parameters. The condition number of FIM indicated information contained in the eigenvalue directions. A rank-deficient FIM with a rank less than the number of parameters pointed towards non-identifiable directions in the eigenspace.

### 3.1 Identifiability analysis revealed redundant kinetic structure and enabled model reduction

The mathematical structure of kinetic rate equations and the identifiability of the model parameters were assessed for the initial 48-parameter model. In the case of non-identifiable parameters, multiple combinations of non-identifiable parameter pairs can produce identical model output, so the optimizer cannot distinguish the true underlying parameter values from any other combination along the same flat direction in parameter space (Raue, Kreutz et al. 2009). Using the Control-condition dataset, 6 of 48 kinetic parameters were found to be non-identifiable.

The magnitude of the first three eigenvalues in Fig 3A (10^−14^ *to* 10^−13^) highlights the non-identifiability of *K*_*dual,Gln*_, *K*_*dual,Glu*_, *K*_*dual,Asn*_, *K*_*dual,Asp*_, *Kd*_*LDH*_ *and α* parameters. All four *K*_*dual*_ parameters were part of the cell growth rate equation, while the other two non-identifiable parameters were part of the death rate equation. The fourth smallest eigenvalue was 10^−2^ after the lowest three, orders of magnitude larger than the first three eigenvalue directions. The non-identifiability artificially inflated the Fisher matrix, leading to a large condition number of 10^23^. Based on the correlation coefficient calculations using *FIM*^−1^, the parameter pairs *K*_*dual,Gln*_ and *K*_*dual,Glu*_, *K*_*dual,Asn*_ and *K*_*dual,Asp*_ and *Kd*_*LDH*_ and *α* were found to be perfectly correlated, with a correlation coefficient of 1 (Fig 3B).

**Fig 3.**
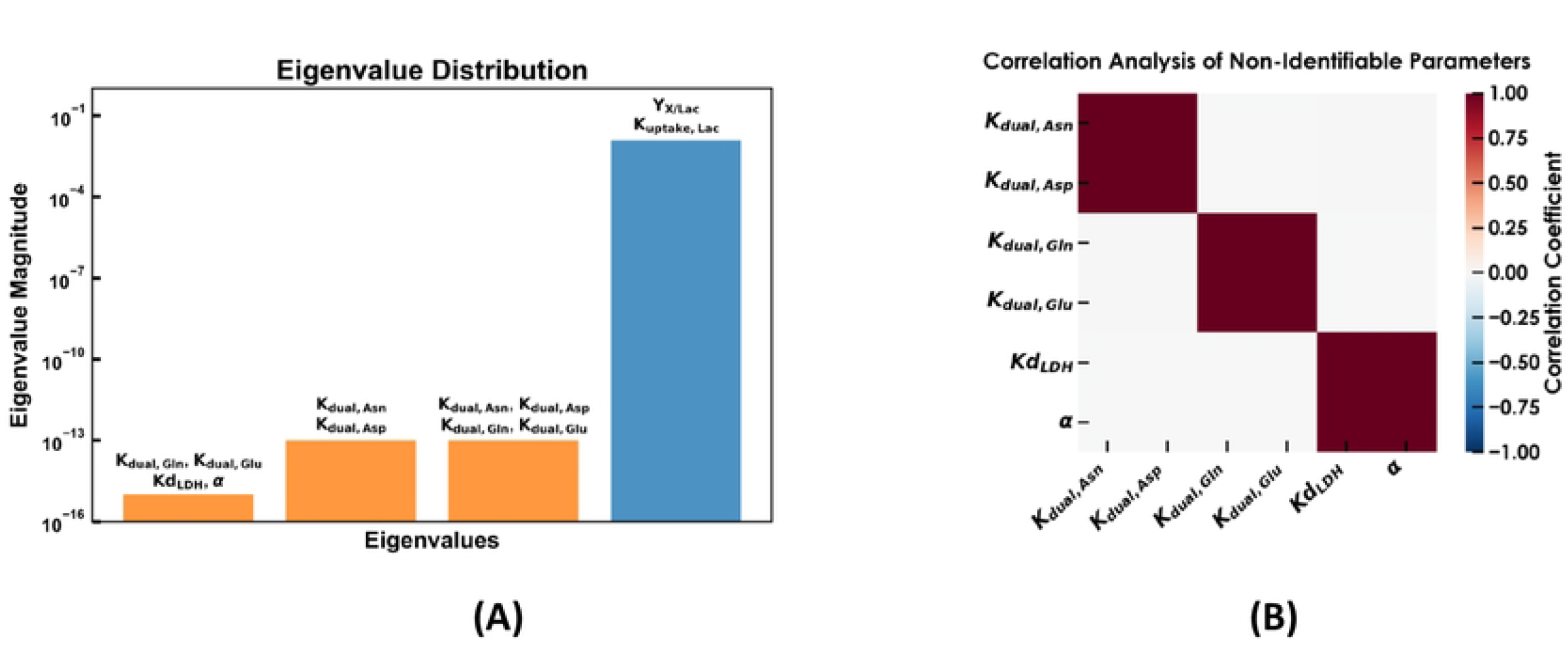
(A) Eigenvalues for non-identifiable parameters are much smaller (∼ - ) than identifiable parameter eigenvalues (≥ -). Eigenvalue analysis informs six non-identifiable parameters. (B) Near-perfect correlation between non-identifiable parameter pairs.

To resolve the non-identifiability of four *K*_*dual*_ parameters, the structure of growth rate (Equations 10 and 11) was assessed.

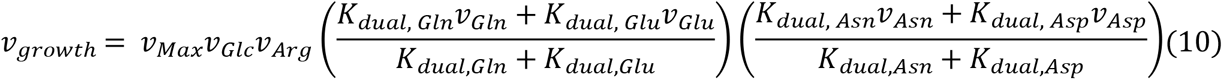

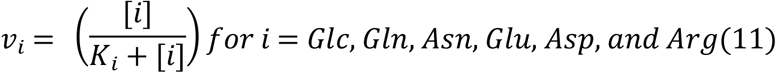

Rewriting Equation 10 by rearranging glutamine and glutamate terms while omitting additional terms for simplicity (Equation 12), only the ratio 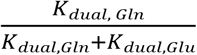 could be estimated based on the mathematical structure of the growth rate equation. Therefore, a new term based on this ratio was defined (Equation 13).

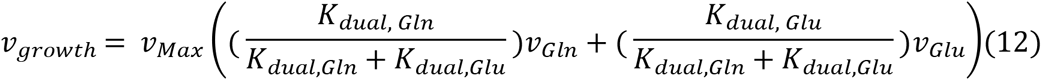

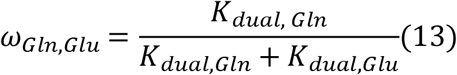

The same algebraic reasoning applies to the asparagine/aspartate pair (*K*_*dual, Asn*_ and *K*_*dual,Asp*_). Both these parameters appear in Equation 10 only as the ratio 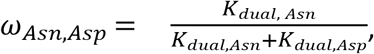, so only their ratio is identifiable and not the individual parameters. The model output was mathematically identical for multiple combinations of (*K*_*dual,Gln*_, *K*_*dual,Glu*_) and ( *K*_*dual,Asn*_, *K*_*dual,Asp*_) pairs. The optimizer was moving along a perfectly flat valley with infinite solutions, resulting in a curvature of 0. Based on the initial estimated values of (*K*_*dual,Gln*_, *K*_*dual,Glu*_) and (*K*_*dual,Asn*_, *K*_*dual,Asp*_) pairs, the values for *ω*_*Gln,Glu*_ and *ω*_*Asn,Asp*_ were obtained (Supplementary file 1). The reparameterization preserved the biological interpretability of the growth term: *ω*_*Gln,Glu*_ = 0 corresponds to growth being entirely glutamate-driven, *ω*_*Gln,Glu*_ = 1 corresponds to growth being entirely glutamine-driven, and intermediate values represent a weighted combination of the two substrates’ contributions to growth. The simplified growth rate equation with *ω*_*Gln,Glu*_ and *ω*_*Asn,Asp*_ is given in Equation 14. The *ω*_*Gln,Glu*_ *and ω*_*Asn,Asp*_ reparameterization is generalizable to any multi-substrate-weighted Monod growth law and preserves the biological interpretation while simplifying the rate equation (Schmidt, Madsen et al. 2008).

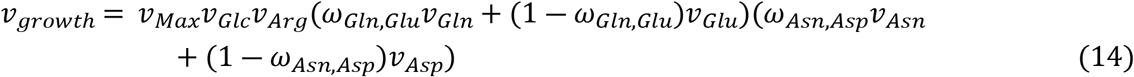

The remaining two non-identifiable parameters *α and K*_*LDH*_ appeared in the death rate equation (Equations 15 and 16) with an eigenvalue of the order 10^−14^. The cell death rate was formulated based on the lactate dehydrogenase (LDH) concentration and viable cell density (VCD). Similar to *K*_*dual*_ terms, *α and K*_*LDH*_ were indistinguishable in the model with a perfect correlation coefficient of 1, since only their ratio (Equation 17) could be estimated. The Control dynamic perfusion condition dataset was used to estimate parameters in the death rate equation because it uniquely captured the cell-death regime following nutrient depletion.

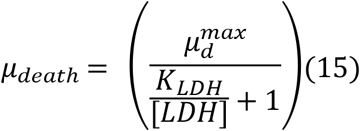

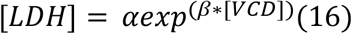

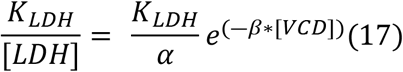

To address the structural issue, the ratio 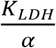 was defined as a new model parameter, *γ* (Equation 18). This simplification maintained the biological dependence of death rate on [LDH] and [VCD] (Equation 19) while resolving the structural discrepancy.

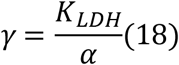

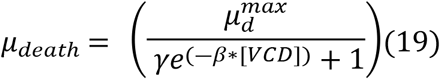

Addressing structural discrepancies in the growth and death rate equations drastically improved the condition number of FIM from 10^23^ *to* 10^7^, thereby removing the non-identifiable directions. The simplification reduced the kinetic model size from 48 to 45 parameters, with a full-rank FIM. Replacing the non-identifiable ratios with single terms for growth and death-rate kinetics did not affect the model fits, as shown in Fig 4. The drastic drop in VCD under dynamic perfusion condition after day 9 was a combined effect of nutrient depletion and filter fouling. The model failed to capture dynamic perfusion after day 9 because filter fouling, a physical rather than metabolic phenomenon, was not accounted for. Two additional parameters *K*_*Glu*_ and 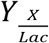 were fixed to their initial estimates to resolve the identifiability issues, bringing the model size to 43 parameters, whose details are provided in Supplementary file 1.

**Fig 4.**
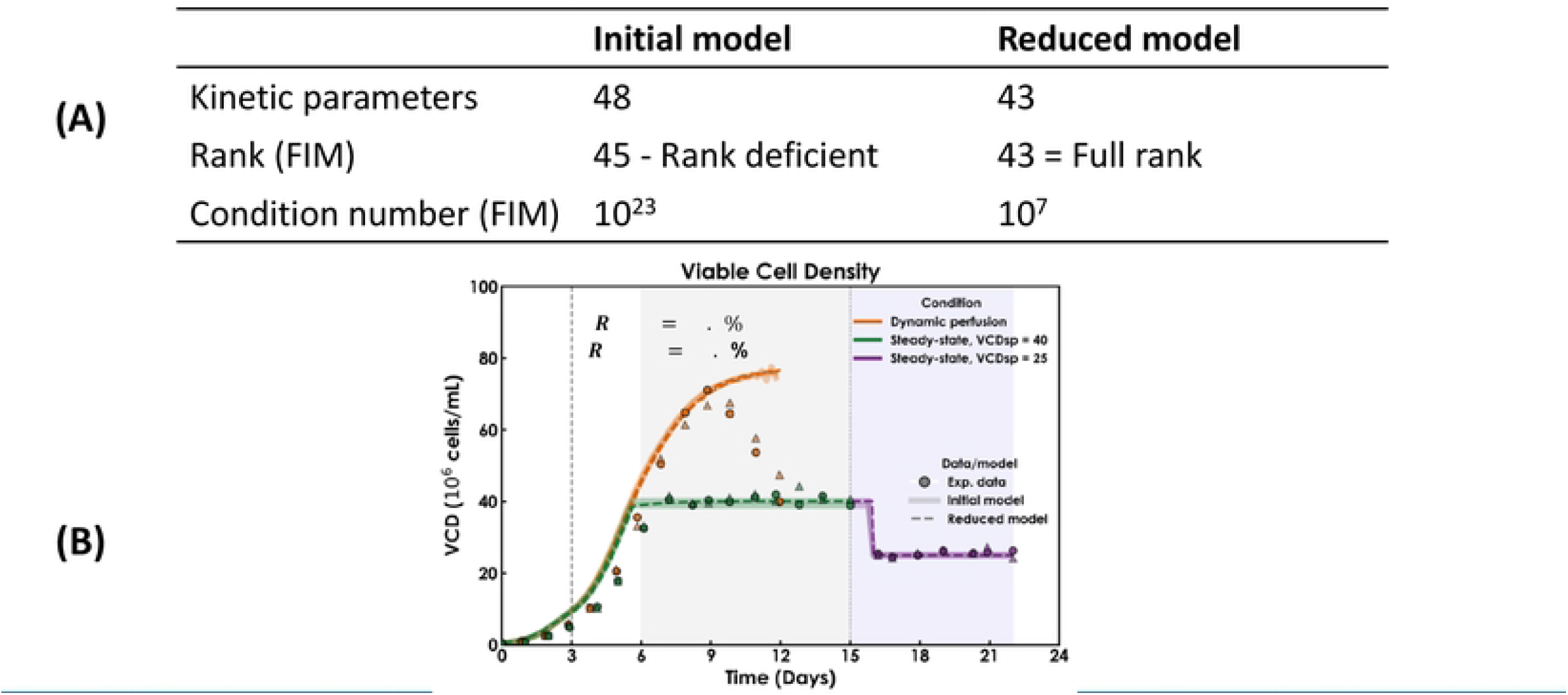
Performance comparison of reduced and initial model. (A) Model performance table. (B) VCD profile for the Control condition. Gray shaded region represents VCD set point of × cells/mL while blue shaded region represents VCD maintained at 25× cells/mL *R* calculations include steady-state operations and omit dynamic perfusion.

For this cell line, glutamine depletion triggered a metabolic switch for lactate, alanine, glutamate, aspartate, serine, and glycine. The published model used binary logic to switch between metabolic phases upon glutamine depletion (Reddy, Malinov et al. 2026). However, binary transitions assume instantaneous, population-wide adaptation, which contradicts biological population heterogeneity. From a mathematical perspective, a binary switch introduces nondifferentiable discontinuities, leading to numerical instability during gradient-based parameter optimization and FIM calculations. Additionally, as observed, a discontinuous binary metabolic switch produces false certainty with a minimal confidence interval, making the uncertainty analysis obsolete (Villaverde, Pathirana et al. 2022).

Lactate metabolism switched from production to consumption once glucose demand for metabolic activities increased (Reddy, Malinov et al. 2026). This shift in lactate metabolism was driven by glutamine concentration, where sufficient glutamine until day 3 led to lactate production, after which lactate metabolism switched to consumption (Equations 20 and 21).

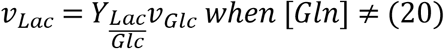

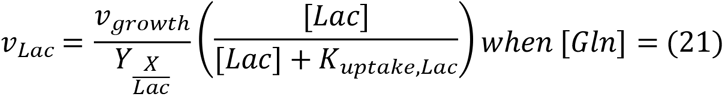

98% correlation and poor identifiability of kinetic parameters *K*_*uptake,Lac*_ *and* 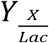 pointed out the discrepancy in the mathematical structure of the lactate rate equation. 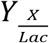 captures the biomass synthesis from lactate and *K*_*uptake,Lac*_ describes lactate consumption for metabolic activities. As both model parameters were required to explain lactate metabolism, they could not be combined or eliminated. To accurately capture the metabolic switch as a function of glutamine concentration, a continuous logistic sigmoid gating function was deployed (Equations 22 and 23). The glutamine gate utilizes an inflection threshold (*Gln*^∗^) and a steepness factor (k). The sigmoid switch is continuous and involves two new parameters k and *Gln*^∗^. k is the rate of metabolic switch and *Gln*^∗^ (intrinsic to the cell line) is an inflection glutamine concentration at which glutamine-dependent metabolic shift is exactly 50% activated. *Gln*^∗^ was estimated as 0.24 mM using lactate flux when the metabolic switch was 50% completed. The estimated value of *Gln*^∗^ is consistent with published values for glutamine affinity constant ranging from 0.2 – 0.6 mM in CHO cells (Ben Yahia, Malphettes et al. 2015).

The term k capturing rate of metabolic transition was estimated by calculating the glutamine concentration transition window over which the metabolism switched for glutamine-concentration-dependent species. For a 10-90% transition width, k was estimated to be 17 *mM*^−1^, assuming the transition in metabolism occurred over a 0 to 0.25 mM glutamine concentration window. Both k and *Gln*^∗^ were excluded from parameter optimization because they govern the structural form of the metabolic switch rather than cell-line-specific metabolic rates.

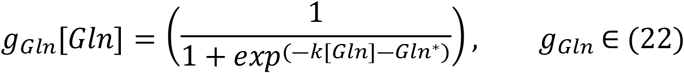

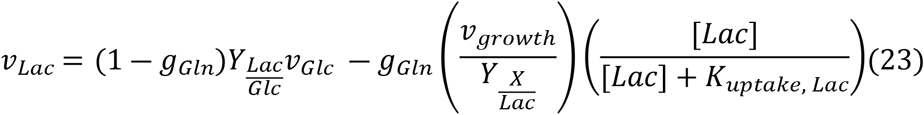

The binary metabolic switch introduced a discontinuity in the mass balance at the glutamine depletion threshold, causing the solver to take excessively small steps near the transition, leading to stiffness. Replacing the binary switch with a continuous sigmoid removed this discontinuity and stiffness. The discontinuity also created a non-differentiable point in the objective function at the metabolic switch time, which destabilized the gradient-based local search within dual annealing. The continuous sigmoid restored a differentiable objective landscape suitable for global optimization. Similar to lactate, alanine, glutamate, serine, aspartate, and glycine exhibited a binary metabolic switch that depended on glutamine concentration and were therefore modeled using the sigmoid function; details are provided in Supplementary file 1. Model reduction significantly improved confidence in the remaining parameters, with CV < 10% for primary stoichiometric yields and maximum specific rates, indicating strong identifiability of central carbon metabolism parameters. Parameters with high uncertainty (CV > 20%) were primarily Monod affinity constants and late-stage death kinetics (Supplementary file 2). Rapid substrate depletion limited data resolution within the death regime, making it difficult to estimate death rate parameters and assess their identifiability.

The reduced dFBA model with 43 kinetic parameters performed similarly to the initial overparametrized model with 48 parameters in explaining the Control condition dataset for all species. MAE and NRMSE values across all 23 species were similar between the initial and reduced models for Control condition (Supplementary file 2). This assessment ensured that simplification yielded a reduced identifiable model with performance similar to the initial over-parametrized model.

### 3.2 Amino-acid supplementation induced a distinct phenotype from the Control medium, which challenged model transferability

The amino acid-supplemented Medium C sustained two cell density steady states at 20 and 30 × 10^6^ cells/mL (Fig 5A). Overall, there was a reduction in specific growth rate for Medium C at both steady states, likely due to reduced amino acid availability (Fig 5B). Along with a reduced specific growth rate, there was a 30% increase in specific productivity at steady state of 30 × 10^6^ cells/mL, creating desirable conditions for perfusion operations (Fig 5C).

**Fig 5.**
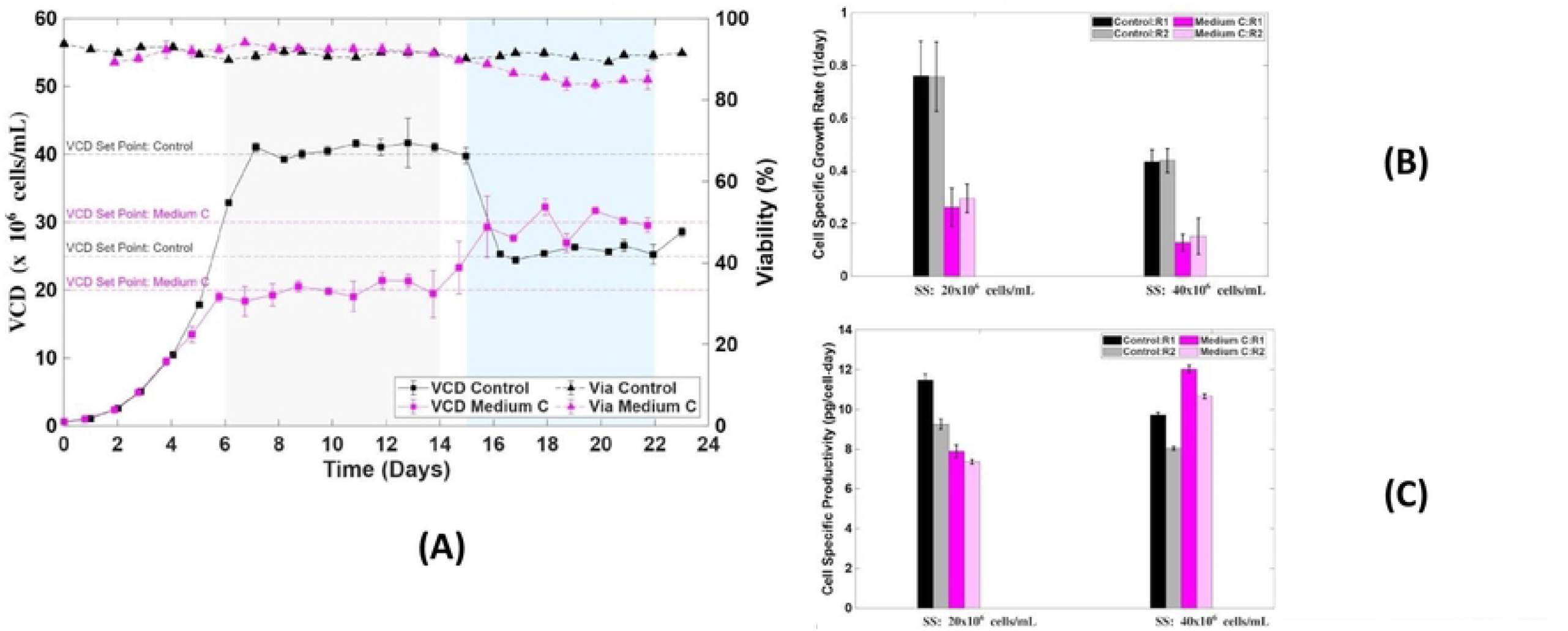
Experimental data for Control and Medium C. (A) VCD and cell viability profiles. Control condition operated at VCD set point of × cells/mLand 25× cells/mL while Medium C was operated at 20× cells/mL and 3 × cells/mL. Point values represent average of two biological replicates and error bars represent standard deviation for each medium. (B) Cell specific growth rates. ( C) Cell specific productivity, which was ∼ 1.3-fold higher productivity for Medium C. Bar height represents average rate for steady state period and error bars represents standard deviation for the period.

Compared to the Control condition, Medium C showed a switch in glutamate and ammonia metabolism from consumption to production at both cell-density steady states. With increased extracellular availability of supplemented amino acids, there was an increase in the uptake rates of tyrosine, tryptophan, and leucine at both steady states.

To test the transferability of the reduced 43-parameter identifiable model, we used it to predict the Medium C phenotype with model parameters estimated using the Control condition. Except for the medium formulation, pH, temperature, and external process parameters were maintained at the same levels as in the Control condition.

The reduced dFBA model trained with the Control condition successfully predicted the VCD (Fig 6A), antibody titer (Fig 6B), and the supplemented amino acids arginine, leucine, tyrosine, and tryptophan (Fig 6C), indicating that the rate equations for these species remained valid under Medium C. However, the model could not capture the shift in glutamate metabolism from consumption for Control to production for Medium C (Fig 7A). Additionally, the model failed to capture lactate production due to metabolic rewiring, thus indicating limitations of the reduced model (Fig 7B). The kinetic model assumed that the rate equations did not change across conditions, only the concentrations that feed into those fixed equations changed. This assumption did not hold for Medium C, where the metabolic flux distribution shifted.

**Fig 6.**
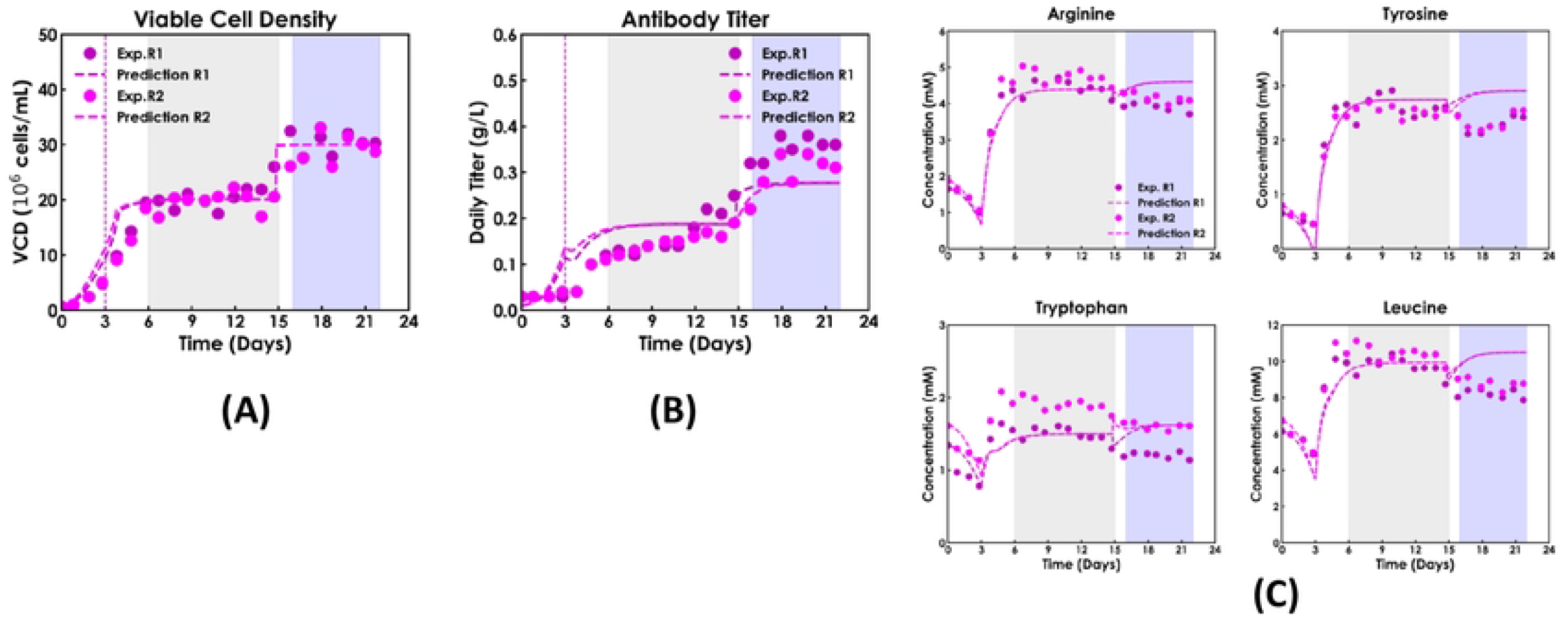
Predictions for Medium C from the reduced model, with kinetic parameters estimated using Control condition only. (A) VCD. (B) Antibody titer. (C) Arginine, leucine, tyrosine, and tryptophan. The two shaded regions represent steady state ranges for the perfusion cultures.

**Fig 7.**
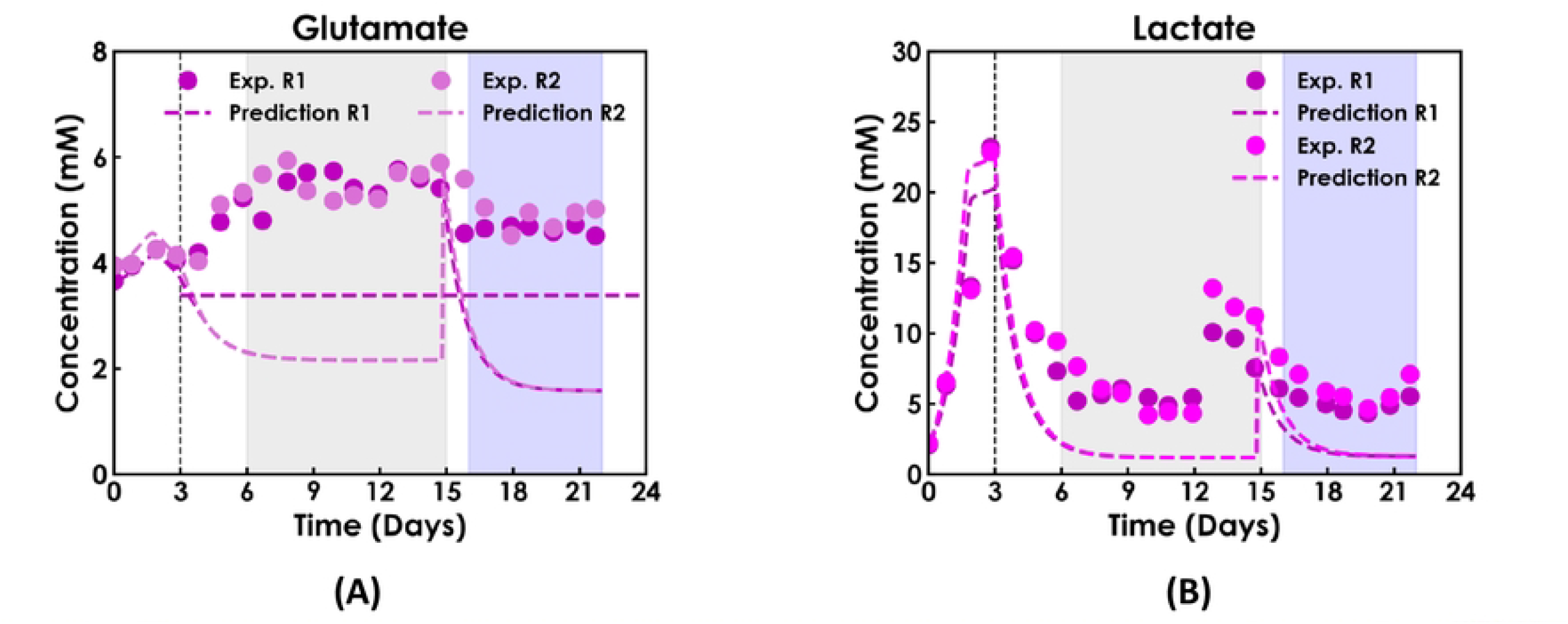
Model predictions in Medium C. (A) Glutamate. (B) Lactate. The model failed to capture reversal of glutamate metabolism and lactate production for Medium C at both steady states. Horizontal dashed line: glutamate concentration in the perfusion medium. The two shaded regions represent steady state ranges for the perfusion cultures.

### 3.3 Parameter-propagated output uncertainty revealed structural inadequacy of the Control medium calibrated kinetic model

The model’s failure to predict glutamate and lactate metabolism for Medium C could reflect either uncertain parameter estimates or structurally inadequate rate equations. To distinguish between these two possibilities, parameter uncertainty was propagated through the model to generate 95% parameter-propagated intervals for Medium C. The Control condition trained parameters were used to calculate parameter-propagated output uncertainty for Medium C using *FIM*^−1^. As shown in Fig 8, experimental trajectories for lactate and glutamate fell completely outside the 95% parameter-propagated intervals, whereas the intervals partially captured the supplemented tryptophan and antibody titer. Combining both biological replicates, 79% of Medium C experimental observations (N = 1058) fell outside the 95% interval, indicating that the Control condition’s calibrated kinetic structure was inadequate to capture lactate production and the reversal of glutamate metabolism in Medium C. The narrow predictive bands, a direct consequence of the low parameter CVs, could not encompass the Medium C experimental trajectories. The narrow bands and 79% points lying outside the intervals were strong evidence of structural inadequacy instead of parameter uncertainty for the prediction failure.

**Fig 8.**
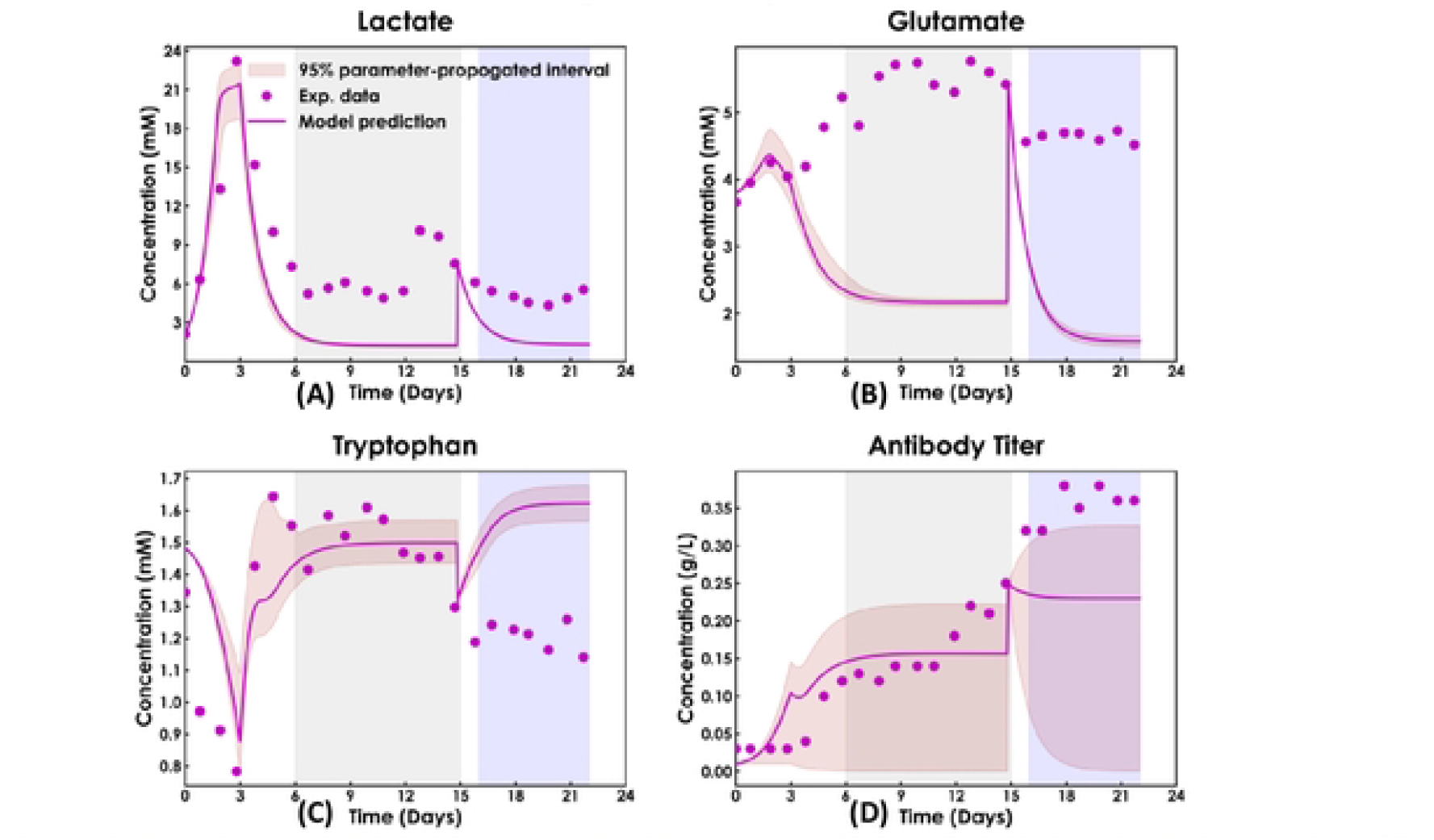
95% parameter-propagated output uncertainty for Medium C. (A) Lactate. (B) Glutamate. (C) Tryptophan. (D) Antibody titer. Glutamate and lactate parameter-propagated intervals did not cover the experimental data while intervals partially captured experimental observations for tryptophan and antibody titer. The two shaded regions represent steady state ranges for the perfusion cultures.

This structural inadequacy of rate equations is consistent with reports that static macroscopic kinetics cannot capture shifts in the global metabolic state without additional constraints on model parameters (Nolan and Lee 2011, Ben Yahia, Malphettes et al. 2015). To investigate the structural failures, MFA was applied using the experimental fluxes as inputs. The exchange reaction fluxes for all 23 species matched with experimental fluxes, confirming that the stoichiometric network reproduced the measured exchange fluxes. Consequently, MFA was used to identify the origin of the kinetic component failure and to guide structural repair for the phenotype observed when using Medium C.

### 3.4 MFA identified the metabolic origin of model failure and guided kinetic model repair

Following the rate equation’s inability to capture the Medium C phenotype and the resulting overconfidence in model predictions due to minimal parameter-propagated interval width, stoichiometric MFA was deployed as a diagnostic tool (Antoniewicz 2013). A flux distribution comparison was carried out across the entire network between the Control and Medium C using MFA. To visualize the flux distribution comparison between the Control and Medium C, a metabolic flux map was created using Escher, with red color indicating backward reactions and teal indicating forward reactions (King, Drager et al. 2015). Only reactions with a significant flux difference between the two media are shown for clarity. Comparing the steady state flux distribution at (40 × 10^6^ cells/mL and 30 × 10^6^ cells/mL) for Control and Medium C, respectively, revealed metabolic rewiring (Fig 9). In the Control condition, ammonia and glutamate were consumed, while these species were produced in Medium C due to extracellular supplementation with leucine, arginine, tyrosine, and tryptophan. Alanine consumption for pyruvate production in the Control condition was reversed in Medium C, resulting in alanine production. Additionally, TCA cycle flux increased in Medium C due to supplementation with essential amino acids.

**Fig 9.**
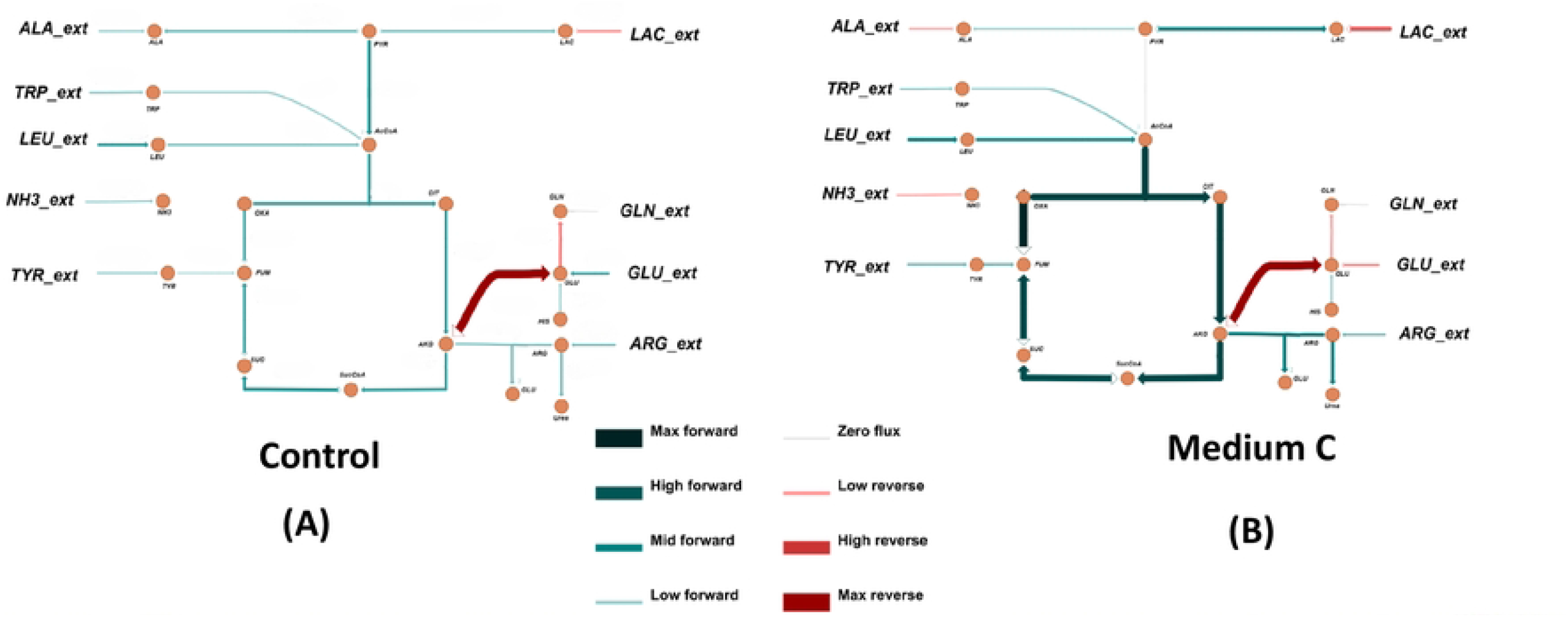
Steady-state metabolic flux comparison for Control (40× cells/mL) and Medium C (30× cells/mL). (A) Control. (B) Medium C. Reversal of ammonia, glutamate and alanine metabolism from Control to Medium C. Increased TCA cycle flux for Medium C was due to supplemented amino acids. Arrow thickness and colour indicate flux magnitude and direction.

MFA identified that glutamate production in Medium C was likely due to the catabolism of supplemented arginine, leucine, and tryptophan. The catabolism of these supplemented amino acids increased the availability of alpha-ketoglutarate, thereby producing glutamate, which was missing from the reduced model’s glutamate rate equation (Plaitakis, Kalef-Ezra et al. 2017). The experimental lactate production observed in Medium C might arise from the catabolism of supplemented amino acids, leading to intracellular NADH accumulation, and thus driving lactate production via the pyruvate-lactate reaction (Mulukutla, Gramer et al. 2012). Again, the reduced model lacked the necessary structure to capture this mechanism.

The detailed comparison of all reaction fluxes was categorized into 3 bins: reactions with a metabolic switch between Control and Medium C, reactions with a magnitude change greater than 50%, and the remaining reactions with a minimal flux change. A detailed comparison of all reactions is provided in Supplementary file 2. For comparison, SS20 refers to steady state at 25 × 10^6^ for Control and 20 × 10^6^ cells/mL for Medium C, and SS40 refers to steady state at 40 × 10^6^ for Control and 30 × 10^6^ cells/mL for Medium C. For both SS20 and SS40, there is a metabolic switch in the alanine, glutamate, and ammonia exchange reactions, as shown in Supplementary file 2. A large change in cell metabolism at SS40 results from increased demand for nutrients and metabolites to support cellular functions, growth, and antibody production.

To ensure that the metabolic flux diagnostics were robust to experimental noise, extracellular flux uncertainties obtained from the two biological replicates for the Control and Medium C were propagated through the stoichiometric matrix via Monte Carlo sampling (N=1000) (Antoniewicz, Kelleher et al. 2006). This analysis revealed statistically significant differences in flux magnitudes between the Control and Medium C, suggesting that improved kinetics are needed to explain the Medium C phenotype. Based on the results, only the reactions showing significant differences between the Control and Medium C were used to guide the structural repair of the kinetic model.

The stoichiometric network provided the missing biological structure, enabling successful capture of the metabolic switch for glutamate and lactate production in Medium C without introducing additional fitted kinetic parameters. Based on the existing kinetic structure, glutamate was consumed only after glutamine depletion on day 3, thereby omitting a term for glutamate production (Equation 24), which is required for Medium C.

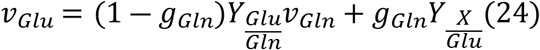

A glutamate production term was added, and the stoichiometric MFA was used to identify the glutamate production sources in Medium C (Equation 25). For Medium C supplemented with arginine, tyrosine, and leucine, they produce glutamate intracellularly (Equations 26-28). With higher availability of arginine, tyrosine, and leucine, cells take up higher amounts of these amino acids, with excess shunted into catabolic pathways, leading to increased TCA flux and production of ammonia and glutamate (Ahn and Antoniewicz 2011).

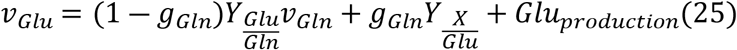

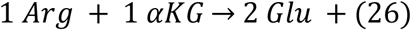

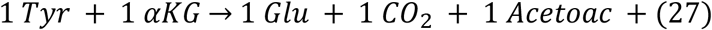

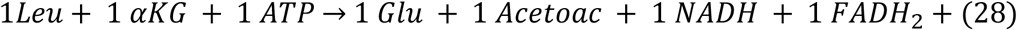

Moving from Control to Medium C, more than 200% increase in flux magnitude was observed in all three reactions across both the steady states (Table 3).

**Table 3.** Metabolic flux comparison for intracellular glutamate-producing reactions. SS20 denotes the steady-state condition at 25×10^6^cells/mL (Control) and 20×10^6^cells/mL (Medium C). SS40 denotes the steady-state condition at 40×10^6^cells/mL (Control) and 30×10^6^cells/mL (Medium C). Tabulated fluxes are averaged over each steady-state period.

| Reaction | Control flux SS20<br>(nmol/$10^6$ cells-day) | Medium C flux SS20<br>(nmol/$10^6$ cells-day) | Control flux SS40<br>(nmol/$10^6$ cells-day) | Medium C flux SS40<br>(nmol/$10^6$ cells-day) |
| --- | --- | --- | --- | --- |
| $\text{Arg} \rightarrow 2\text{Glu}$ | 41.6 | 209.3 | 50.6 | 234.6 |
| $\text{Tyr} \rightarrow \text{Glu}$ | 1.1 | 53.1 | 2.3 | 54 |
| $\text{Leu} \rightarrow \text{Glu}$ | 35.7 | 139.8 | 43.3 | 144.4 |

MFA indicated catabolism of arginine, leucine, and tyrosine as glutamate production sources in Medium C. Each amino acid’s catabolic contribution to glutamate production flux was modeled as directly proportional to its concentration in excess of the Control-condition steady state baseline (Equation 29). Their catabolic contribution was assumed to be zero when the concentration was the same as the Control condition’s baseline, and to increase linearly as the concentration rose above it. This mathematical structure is inspired by Michaelis-Menten kinetics operating in the linear region, where the substrate concentration is below the saturation point (Saa and Nielsen 2017).

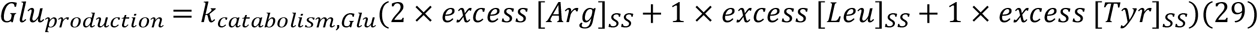

In Equation 29, the coefficients are the stoichiometric yield of glutamate per mole of each amino acid, and *k*_*catabolism,Glu*_ is a single rate constant common to all three amino acids. The first-order rate constant *k*_*catabolism,Glu*_ was calculated for each of the two steady states independently. To estimate *k*_*catabolism,Glu*_, MFA-derived fluxes were used along with excess amino acid concentrations as the driving force for catabolism. By exchange convention, positive flux denotes net consumption while negative flux denotes production. The detailed calculations for *k*_*catabolism,Glu*_ are provided in Supplementary file 1. The *k*_*catabolism,Glu*_ value of 9.5, calculated from the SS40 data, was used in the model’s glutamate rate equation. The SS20 MFA flux and concentrations led to *k*_*catabolism,Glu*_ value of 12.9, which was higher than the SS40 value. *k*_*catabolism,Glu*_ values calculated independently for the two steady states differed, indicating that the first-order approximation captures the primary concentration-dependent behavior but not the full steady-state dependence. With the *k*_*catabolism,Glu*_ value estimated using the SS40 data, the residual glutamate concentration for Medium C was sufficiently captured for 20 × 10^6^ cells/mL steady state but overpredicted for 30 × 10^6^ cells/mL, highlighting a need for a concentration- or CSPR-dependent rate law (Fig 10A).

**Fig 10.**
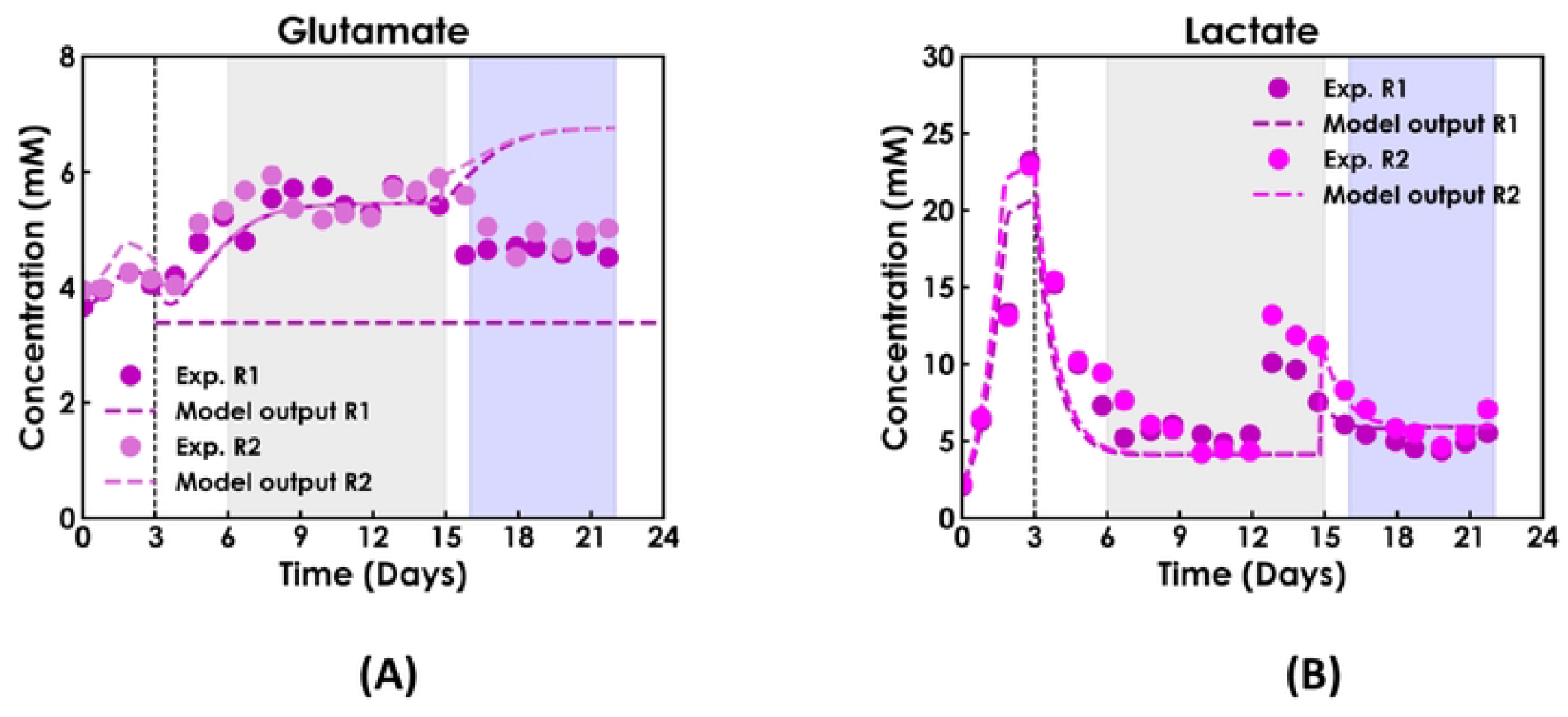
Model output profiles for (A) Glutamate. (B) Lactate. MFA-derived glutamate and lactate production terms reproduced the lactate profile in Medium C, with glutamate overpredicted at higher-density steady state. The two shaded regions represent steady state ranges for the perfusion cultures.

Similarly, the failure to predict lactate production in Medium C at both steady states indicated that a term was missing in the lactate rate equation in Equation 23. The current lactate rate equation produces lactate only when glutamine is available, then switches to lactate consumption after day 3 following glutamine depletion. However, lactate production was observed experimentally in Medium C. Catabolism of arginine, leucine, and tryptophan (Equations 26, 28, and 30) generates excess NADH. LDH regenerates *NAD*^+^ from this surplus NADH by reducing pyruvate to lactate (Equation 31). This NADH-driven flux through LDH, rather than the amino acids themselves, is the mechanistic link between their catabolism and increased lactate production (Mulukutla, Gramer et al. 2012). Intracellular NADH accumulation flux was obtained using the MFA-derived fluxes from the supplemented amino acids, arginine, tryptophan, and leucine, with more than a 200% increase in reaction fluxes in Medium C compared to the Control condition (Table 4).

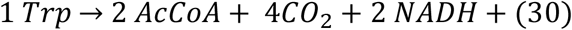

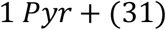

**Table 4.**
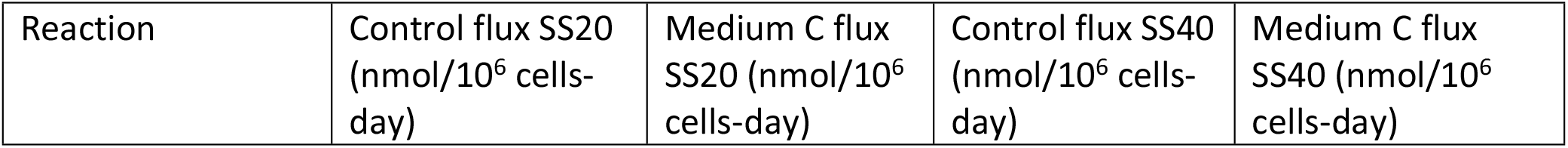
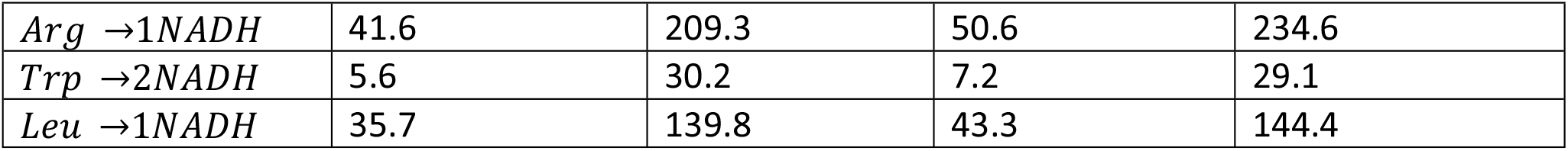
Metabolic flux comparison for intracellular NADH-producing reactions. SS20 denotes the steady-state condition at 25×10^6^cells/mL (Control) and 20×10^6^cells/mL (Medium C). SS40 denotes the steady-state condition at 40×10^6^cells/mL (Control) and 30×10^6^cells/mL (Medium C). Tabulated fluxes are averaged over each steady-state period.

Similar to glutamate rate modification, a lactate production term was added to the lactate rate equation (Equation 32). The three supplemented amino acids, arginine, tryptophan, and leucine, were selected as catabolic sources for NADH accumulation, thereby driving the production of lactate from pyruvate. Their catabolic contribution to lactate production flux follows the same excess-concentration structure (Equation 33).

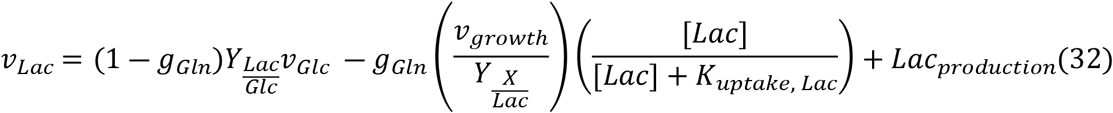

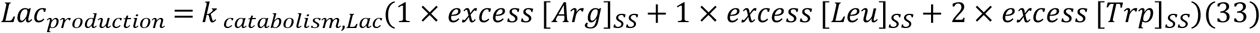

The coefficients represent the stoichiometric yield of lactate per mole of each amino acid and *k*_*catabolism,Lac*_ is a single rate constant common to all three amino acids in Equation 33. The derivation for *k* _*catabolism,Lac*_ is provided in Supplementary file 1. The *k*_*catabolism,Lac*_ value of 10.7 (calculated from the SS40 data) was used in the model’s lactate rate equation. Using the SS20 data, *k*_*catabolism,Lac*_was estimated to be 18.4, higher than the rate constant obtained from the SS40 data. Lactate production in Medium C is correctly captured with the *k*_*catabolism,Lac*_ value estimated using the SS40 data for both steady states (Fig 10B).

The model improvement was quantified using the NRMSE and MAE scores. While glutamate and lactate NRMSE values improved by 64% and 36%, respectively, in Medium C, three species – alanine, glutamine, and valine showed NRMSE increases exceeding 10% (Supplementary file 2). NRMSE values for these species changed only during identifiability-driven model reduction in section 3.1. The values had negligible changes during MFA-guided structural repair, confirming that MFA-guided repair was localized to glutamate and lactate. The remaining species showed NRMSE changes within ±10%, and overall model performance across 23 species was maintained. Additionally, the MAE improved by 71% for glutamate and 57% for lactate.

### 3.5 Complementary metabolic conditions confirmed identifiability of the repaired model and improved parameter precision

With different cell metabolisms for Control and Medium C, complementary information with different sensitivities for model parameters was obtained, further reducing the uncertainty in model parameters and outputs when combining information from both media (Franceschini and Macchietto 2008). To highlight the benefit of Medium C in addition to the Control condition, an identifiability analysis was performed using the following 3 dataset: the Control condition, the Medium C condition, and a single dataset combining the Control and Medium C conditions.

*k*_*catabolism,Lac*_ and *k*_*catabolism,Glu*_ were added to the kinetic parameter list to quantify model improvement using FIM-based metrics but were not optimized. The Control condition provided estimates of growth-linked yield parameters but showed minimal sensitivity for MFA-guided *k*_*catabolism,Lac*_ and *k*_*catabolism,Glu*_, because the supplemented amino acid concentrations did not exceed the Control threshold. Combining experimental information from Medium C with the Control condition provided orthogonal information, informing parameter directions that the Control condition alone could not - providing estimates for *k*_*catabolism,Lac*_ and *k*_*catabolism,Glu*_. With combined Control and Medium C data, the condition number of FIM reduced by 16% compared to the Control condition alone, and the rank of FIM matched the number of parameters. The 16% reduction in condition number understates the information gain, as the condition number reflects only the ratio of extreme eigenvalues and was dominated by persistently non-identifiable *K*_*uptake,Lac*_. Combining Control and Medium C data drastically reduced the CV values for *k*_*catabolism,Lac*_ and *k*_*catabolism,Glu*_ from 36.5% and 28.0% in the Control dataset to 4.4% and 1.9%, respectively. The information gain in the eigenvalue direction associated with *k*_*catabolism,Lac*_ and *k*_*catabolism,Glu*_ improved by approximately two orders of magnitude (from 36.5% and 28.0% to <5%, respectively), reflecting information gain that Control data alone cannot improve.

Of the 20 growth-linked amino acid yield parameters, 16 had CV values lower than 10% using the combined data, consistent with the current mathematical structure for amino acid metabolism and providing sufficient confidence in the parameter estimates (Supplementary file 2). The assessment of pairwise correlations between parameters revealed no structural issues, as none of the correlation values exceeded 80%.

## 4. Discussion

When assessing the prediction capabilities of models, typically only the mean performance is evaluated, while confidence in those predictions is ignored (Anane, López C et al. 2019, Bayer, Duerkop et al. 2023). *FIM*^−1^ calculation is computationally cheaper and provides a lower bound on model parameter estimates, which can be used to generate model output uncertainties (Villaverde, Raimundez et al. 2023). A more detailed kinetic model is not necessarily more informative, as demonstrated through the model reduction exercise, in which 43 parameters were sufficient to capture cell metabolism while preserving biological interpretation. Fekih-Salem applied similar model-reduction approaches to reduce model size while preserving mechanisms (Fekih-Salem, Dewasme et al. 2019).

As demonstrated, a reduced and well-identifiable model is not definitively transferable, even when the Control-condition-trained kinetic parameters were identifiable and were generating excellent model fits. However, under the metabolic perturbation in Medium C, the reduced model struggled to predict glutamate and lactate. This finding demonstrated that parameter precision and structural adequacy are different dimensions and need to be assessed individually. The stoichiometric network successfully localized the metabolic origin of the model’s structural failure. It supplied repair terms directly from flux data, avoiding the common alternative of incorporating each new phenotype as an additional empirical parameter (Nolan and Lee 2011).

Based on the Cramer-Rao bound, *FIM*^−1^ provides a lower bound on model parameter uncertainties, indicating the actual uncertainties might be larger. Additionally, local sensitivity-based FIM analysis is sensitive to initial parameter estimates. It may require multiple iterations to reduce parameter uncertainties when the initial estimates are far from the true values (White, Tolman et al. 2016, Wieland, Hauber et al. 2021). When performing the FIM analysis, these limitations need to be underlined. Despite its limitations, FIM analysis has been shown to reduce model size and parameter uncertainties effectively (Fekih-Salem, Dewasme et al. 2019, Wang and Dowling 2022). A major benefit of FIM is the computational efficiency compared to Sobol or Monte Carlo sampling, which requires model simulations for each parameter combination, significantly increasing the computational load, especially for large-scale metabolic models (Sobol’ 2001).

FIM-based confidence intervals assume a Gaussian approximation to the likelihood surface (Raue, Kreutz et al. 2009). Some of the model parameters exhibited strongly asymmetric 95% confidence intervals, *γ* with a value of 371.9 and confidence interval ([59.6, 2319.9]), *K*_*uptake,Lac*_ with a value of 9.8 and confidence interval ([2.0, 46.9]), violating the Gaussian surface assumption. For these parameters, FIM-derived CV underestimated true uncertainty, and the profile likelihood method would provide more reliable confidence intervals (Raue, Kreutz et al. 2009). However, computational tractability makes FIM an appropriate first-order screening tool (Villaverde, Raimundez et al. 2023). For most of the kinetic parameters in the model, the confidence intervals were symmetric, consistent with the FIM Gaussian assumption, thereby making the eigen analysis and CV values reliable (Supplementary file 2).

Two parameters remained poorly identified in the final reduced and repaired model, *K*_*uptake,Lac*_ and *γ*. High uncertainty in *K*_*uptake,Lac*_ was due to the minimal sensitivity of this parameter, since neither of the media led to a saturation regime for lactate concentration to estimate *K*_*uptake,Lac*_ confidently. High uncertainty in the death rate parameter *γ* arose since dynamic perfusion condition provided limited information about the cell death regime, which was insufficient to estimate *γ* confidently. To estimate *γ*, experiments require high-cell-density conditions with a sustained high death rate. Based on the optimality conditions, subsequent experiments can be designed and screened using alphabetic optimality criteria to estimate these parameters (Pankajakshan, Waldron et al. 2019, Shahmohammadi and McAuley 2019, Gevertz and Kareva 2024).

Together, these results demonstrate that a reduced, well-identifiable kinetic model is not automatically transferable across different phenotypes. The stoichiometric MFA offered a principled route to diagnose and repair structure without expanding the model size. The integrated identifiability MFA approach can be adapted for cell metabolic models to reduce, repair, and successfully transfer the model across different phenotypes.

## 5. Code and Data Availability

The experimental data were generated within the AMBIC consortium, and the model code is part of intellectual property filed at the University of Delaware. Their public release is therefore subject to consortium data-sharing agreements and institutional review, which are in progress. All equations, parameter values, and processed data underlying every figure and table are provided in main text and Supplementary Information, which is sufficient to reimplement the model and reproduce the reported results. We will provide the material to the editors and reviewers on request during peer review, and we intend to deposit the code and data in a public repository upon publication, to the extent permitted by these agreements.

## 7. Nomenclature

**Model, statistical and process terms**

CI: Confidence interval
CSPR: Cell-specific perfusion rate
CV: Coefficient of variation
dFBA: Dynamic flux balance analysis
FIM: Fisher information matrix
MAE: Mean absolute error
MFA: Metabolic flux analysis
NRMSE: Normalized root mean squared error

**Metabolites and biochemical species**

AcCoA: Acetyl-coenzyme A
Amm: Ammonia
Arg: Arginine Asn - Asparagine
Asp: Aspartic acid
ATP: Adenosine Triphosphate
Cys: Cysteine
Fum: Fumarate
Glc: Glucose
Gln: Glutamine
Glu: Glutamic acid
G6P: Glucose-6-Phosphate
Gly: Glycine
His: Histidine
Ile: Isoleucine
Lac: Lactate
LDH: Lactate dehydrogenase
Leu: Leucine
mAb: Monoclonal antibody
Met: Methionine
NADH: Nicotinamide adenine dinucleotide
Phe: Phenylalanine
Pyr: Pyruvate
Trp: Tryptophan
Tyr: Tyrosine
Ser: Serine
Val: Valine
VCD: Viable cell density
X: Biomass
*αKG*: Alpha ketoglutarate

## 8. Appendix A: Kinetic equations

**Table A1:**
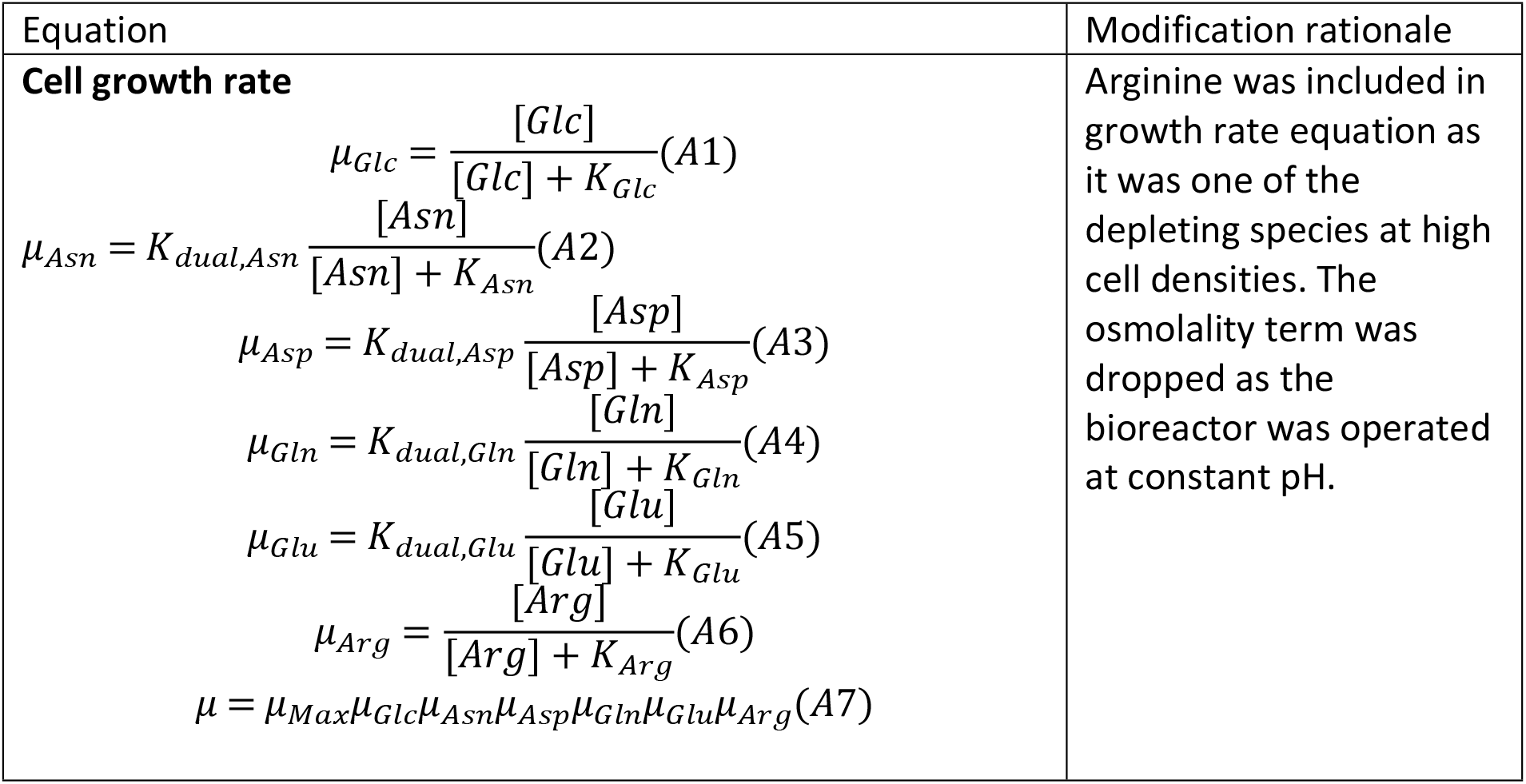
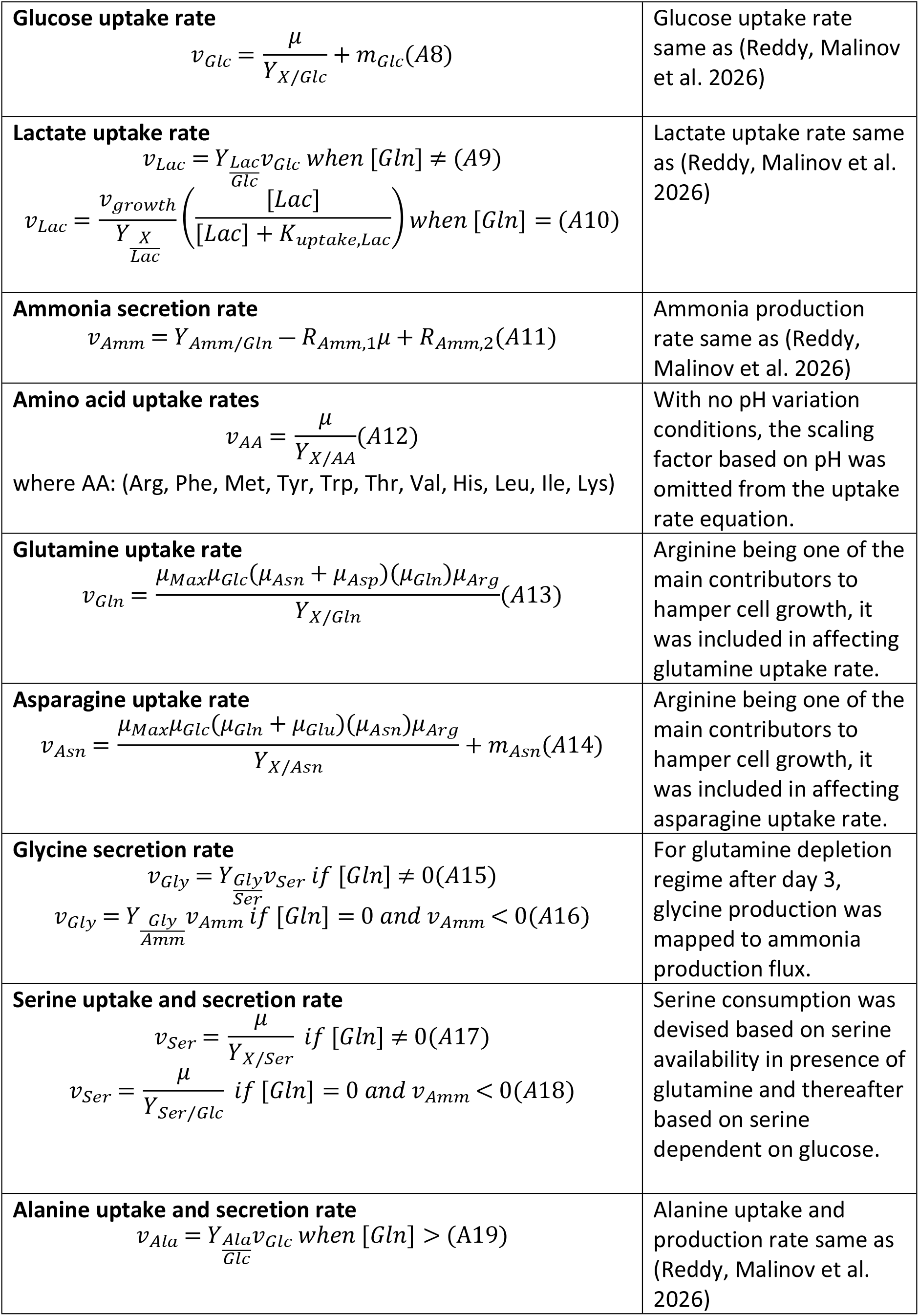
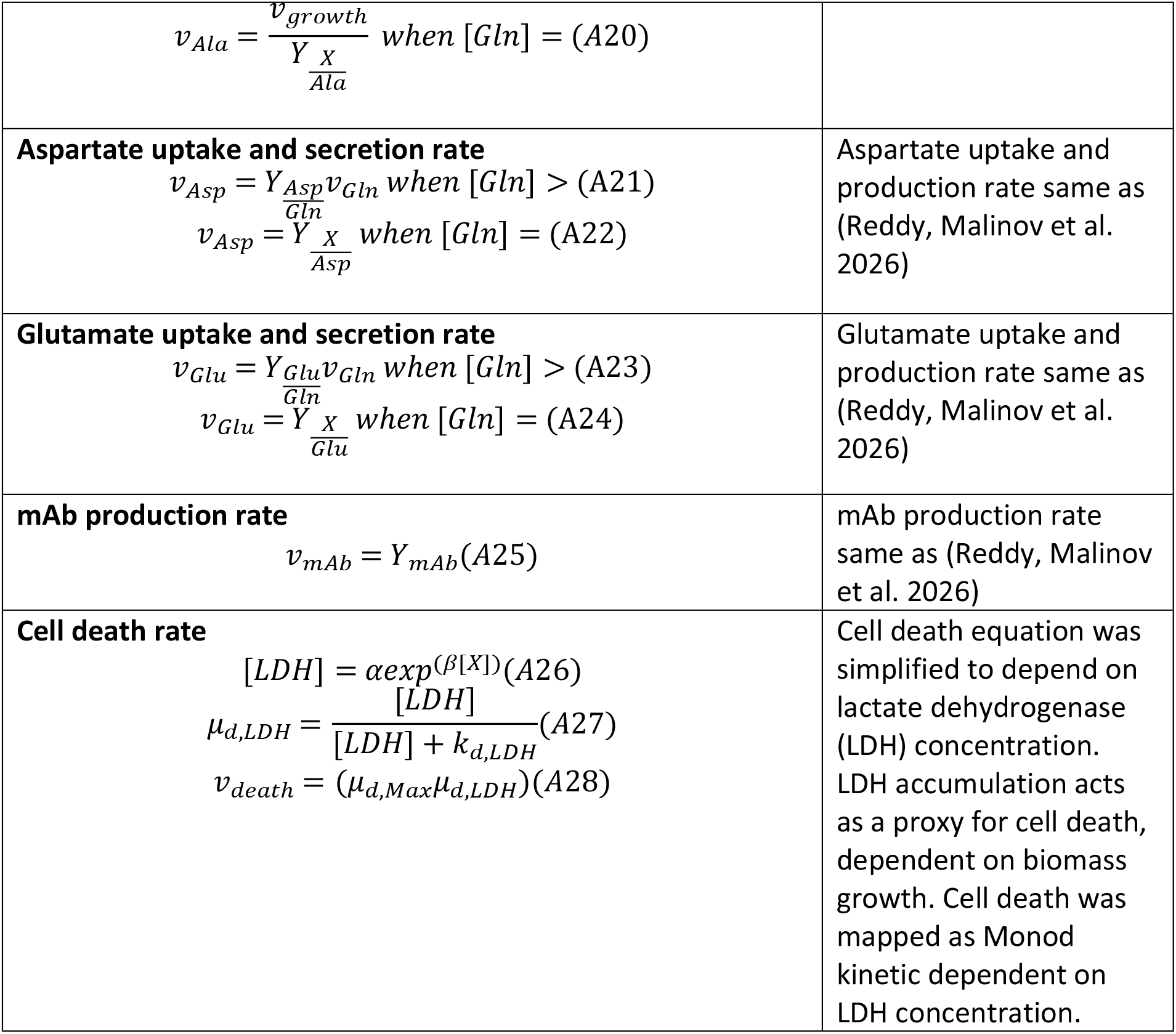
Same kinetic equations as (Reddy, Malinov et al. 2026) were used with the below mentioned modifications.

## 9. Appendix B: Reaction network

**Table B1:** Additional and modified reactions in reaction network for metabolic flux analysis. Same metabolic reaction network as (Reddy, Malinov et al. 2026) was used except for the below mentioned modifications.

| Reaction number | Reaction |
| --- | --- |
| B1 | $Arg + \alpha KG \rightarrow 2Glu + 1NADH$ |
| B2 | $Gly \rightarrow CO_2 + NH_3 + NADH$ |
| B3 | 0.01259 *Ala* + 0.01025 *Arg* + 0.00705 *Asn* + 0.01509 *Asp*<br>+ 0.00312 *Cys* + 0.01505 *Gln* + 0.00457 *Glu*<br>+ 0.01645 *Gly* + 0.00308 *His* + 0.00603 *Ile*<br>+ 0.01368 *Leu* + 0.01154 *Lys* + 0.00308 *Met*<br>+ 0.00851 *Phe* + 0.01123 *Ser* + 0.00855 *Thr*<br>+ 0.00223 *Trp* + 0.00467 *Tyr* + 0.00834 *Val*<br>+ 0.6138 *ATP* + 0.0025 *NH<sub>3</sub>* + 0.07733 *NADH*<br>+ 0.05304 *AcCoA* + 0.01716 *G6P* → 0.00812 *CO<sub>2</sub>*<br>+ 0.0039 *Fum* + 1.0 *Biomass* |

## 10. Contributions

Shivam Barodiya: Conceptualized, Data curation, Investigation, Formal analysis, Visualization,

Writing – original draft preparation, Writing – review and editing.

Nikola Malinov: Methodology, Software.

Srikanth Rapala: Acquisition, Data curation

Marianthi Ierapetritou: Conceptualization, Resources, Project administration, Writing – review and editing.

Sarah Harcum: Project administration, Writing - editing.

